# Dynamics of cutaneous infection of a *Bacillus cereus* biovar *anthracis* strain

**DOI:** 10.64898/2026.09.28.754931

**Authors:** Annabelle Garnier, Clémence Rougeaux

## Abstract

*Bacillus cereus* biovar *anthracis* (Bcbva) strains, which cause fatal anthrax-like disease, have been isolated from animals in Central and West Africa. Like *B. anthracis*, Bcbva produces a poly-γ-D-glutamic acid (PDGA) capsule, as well as edema factor (EF), lethal factor (LF) and protective antigen (PA). Bcbva also possesses a hyaluronic acid capsule that is not expressed by *B. anthracis*. The pathogenesis of Bcbva has recently been characterized in inhalational infection models, in rabbits and guinea pigs. Here, we characterized the kinetics of Bcbva infection and toxin production in a murine model of cutaneous infection using the Bcbva CA strain isolated in Cameroon. We found that spore germination occurred rapidly at the inoculation site, followed by bacterial dissemination to the draining cervical lymph nodes. Germination was associated with rapid production and diffusion of LF and EF. Bacteria rapidly disseminated to lymphoid organs and other compartments, with distal germination observed 18 hours post-infection. Infected mice developed septicemia within 24 hours, accompanied by high circulating LF levels and signs of cardiac and renal injury. These findings may contribute to the fulminant course of Bcbva infection observed in animals and highlight its potential risk to humans.

## 1. Introduction

*Bacillus anthracis*, the agent of anthrax, is a Gram-positive spore-forming bacteria found worldwide. It is genetically close to *B. cereus*, both bacteria belonging to the *B. cereus* group [1]. Over the past twenty years, emerging atypical *B. cereus* strains have been described, responsible for anthrax-like diseases. Thus, *Bacillus cereus* biovar *anthracis* (Bcbva) strains have been isolated from wild animals in tropical rainforests in Cameroon, Côte d’Ivoire and the Central African Republic, as well as from domestic animals in the Democratic Republic of the Congo [2–6]. These atypical *B. cereus* strains are suspected to be major agents of infection in wildlife [2,7], affecting various animal species including chimpanzees, duikers, goats, elephants and great apes [2,4,6,8,9]. However, no confirmed human cases of anthrax-like disease have been reported to date, despite the detection of antibodies against a Bcbva-specific protein (pX02-60) in more than 10 % of a rural human population tested in Côte d’Ivoire [10,11].

Bcbva isolates combine phenotypic characteristics of a) *B. anthracis* (the lack of beta-hemolytic activity and an inactive pleiotropic regulator PlcR –with mutations differing between Bcbva and *B. anthracis-*) and b) *B. cereus* (motility, resistance to gamma phage, and in some isolates, resistance to penicillin G) [5,6,12]. They harbor a chromosomal backgroung derived from a non-*B. anthracis* member of the *B. cereus* group, and belong to a distinct clade from that of *B. anthracis* [6]. However, they carry two plasmids, pBCXO1 and pBCXO2, which are highly similar genetically to the plasmids of *B. anthracis*, pXO1 and pXO2 [4–6,13]. Like plasmids pXO1 and pXO2, plasmid pBCXO1 encodes three toxin components, protective antigen (PA), edema factor (EF), and lethal factor (LF) whereas pBCXO2 encodes a poly-gamma-D-glutamic acid (PDGA) capsule. Bcbva also produces a hyaluronic acid (HA) capsule, encoded by pBCXO1, which is not expressed by *B. anthracis* [14]. The HA capsule contributes to the virulence of Bcbva, which is similar to that of the fully virulent *B. anthracis* 9602 strain in guinea pig and murine models of cutaneous and inhalational disease [15].

Disease progression associated with Bcbva strains isolated in Côte d’Ivoire and Cameroon has recently been characterized in guinea pig and rabbit models of inhalational disease [16,17]. In the study of Ferris et *al*. [16], the authors demonstrated that Bcbva isolates in Côte d’Ivoire and Cameroon cause a disease similar to that caused by *B. anthracis* in terms of virulence, bacterial load and virulence factor expression, including PA and PDGA production. In the study of Jiranantasak et al. [17], the Bcbva strain isolated in Côte d’Ivoire was shown to be more virulent than the *B. anthracis* Ames strain in guinea pigs and *Galleria mellonella* larvae. Both studies focused on the inhalational route of infection, which leads for *B. anthracis* to the most fulminant and lethal form of anthrax.

The route of entry of Bcbva remains unclear. Therefore, gastroinstestinal and cutaenous routes of infection must not be neglected. Tropical rainforest flies may represent a potential source of contamination, either by contaminating vegetation and the environment [8,18,19], where Bcbva may form biofilms [20], or by biting animals as has been suspected and demonstrated for *B. anthracis* with hematophagous insects and flies [21–26]. Based on results obtained with laboratory *B. anthracis* strains, it was estimated that a single blow fly could contaminate leaves with up to 8.62 x 10⁵ spores per day, with a carcass that could potentially harbor thousands of flies [19]. Cultural practices, such as cutaneous scarification or handling of contaminated bushmeat may also represent risk factors for Bcbva entry through skin wounds [27–29], as observed during recent outbreaks of human cutaneous anthrax and with other pathogens [30–33]. The zoonotic risks associated with the bushmeat trade were recently analyzed in the Democratic Republic of the Congo [27]. Bcbva was detected in 0.5% of all animal samples collected at the market; these animals were probably found dead. Handling contaminated meat poses a potential risk to hunters, transporters, butchers, sellers and buyers and human activities may facilitate the spread of Bcbva, as has been described for *B. anthracis* [34]. Therefore, we consider it important to investigate the cutaneous route of infection.

Here, we characterized the toxi-infection associated with the fully virulent Bcbva strain CA, isolated in Cameroon, using a murine model of cutaneous disease, to decipher the associated pathogenicity and to assess the potential risk of a human cutaneous form of anthrax-like disease. The virulence of the Bcbva strain CA was first verified in *Galleria mellonella* (*G. mellonella*) larvae, to reduce the use of vertebrate animals. Afterwards, in immunocompetent mice, we highlighted rapid bacterial germination in the injected ear, and the early presence of bacilli in the cervical lymph nodes draining the inoculated ear. Bacterial germination was associated with the rapid production and diffusion of LF and EF. We showed that mice developed septicemia within a day post-infection as a consequence of the rapid progression of infection. An early increase in cardiac troponin, a biomarker of cardiac lesion, and an increase of creatinine, a biomarker of renal function were also observed. These combined events may explain the fulminant and severe systemic effect of cutaneous infection with the bcbva strain CA.

## 2. Materials and Methods

### Chemicals and Reagents

Recombinant edema factor (EF) and lethal factor (LF) from *Bacillus anthracis* were purchased from CliniSciences (Nanterre, France). Adenosine 5′-triphosphate (ATP) disodium salt hydrate, sodium periodate, rhamnose, were from Sigma-Aldrich (St. Louis, MO). Calmodulin human recombinant (rCaM) was from Enzo Life Sciences (Villeurbanne, France). Pre-coated mouse anti-rabbit IgG plates, cAMP acetylcholinesterase (AChE) enzymatic tracer, Ellman’s reagent and acetic anhydride used in the EIA studies were from Bertin Health & Life Sciences (Montigny-Le-Bretonneux, France). Casein and streptavidin poly-HRP were from Thermo Fisher Scientific (France).

### Ethics statement

All experimental procedures were performed in compliance with the European Directive 2010/63/EU and French legislation (edict 2013-118) and after approval by the Institutional Animal Care and Research Advisory Committee of the French Army Health Service (authorization 511985, July 9 2024). The animals (six per cage) were acclimated for at least a 7-day period before the beginning of the experiments in an environment maintained between 20 and 22°C and a 12-h dark/light cycle. Food and tap water were given *ad libitum*. The animals were controlled daily.

### Virulence assay of CA

The Bcbva strain CA was isolated in Cameroon (kindly provided by Silke Klee, Robert Koch-Institute, Centre for Biological Threats and Special Pathogens, ZBS 2: Highly Pathogenic Microorganisms). CA produces two capsules (a PDGA capsule and a hyaluronic acid capsule) and two toxins, composed by the protective antigen (PA), edema factor (EF) and lethal factor (LF) [15]. All experiments were conducted in biosafety level 3 (BSL3) laboratory. Before mice infection, we verified the virulence of Bcbva CA used in the study in larvae *Galleria mellonella*.

*Galleria mellonella* larvae weighting between 100-200 mg were purchased from INRAE (Jouy-en-Josas, France). Bcbva CA spores were prepared at the desired concentration. Larvae were briefly placed at +4°C to reduce mobility [17]. 10 μL of spores was injected in the uppermost proleg of immobile larvae, as previously described [35], using an infusion pump KDS 100Y (Delta Labo, France), a 1mL syringe (UGAP, France) and a Venofix® A 27G (Dominique Dutscher, France). Injected larvae with 10 μL of phosphate-buffered saline (PBS) (VWR, France) were used as negative control. Larvae were placed in Petri dishes and incubated at 37°C in the dark. Larvae were monitored every day. Surviving larvae after 10 days of monitoring and dead larvae were placed at -20°C before their elimination. Each experiment was carried out in triplicate, with 10 to 12 larvae per condition.

### LF and EF production *in vitro*

Spore germination of Bcbva CA and *B. anthracis* 9602-*lux* was induced in Brain Heart Infusion (BHI) broth (Sigma, France), at 37°C, 5 % CO_2_. After 10 minutes of incubation in BHI broth, the culture was pelleted by centrifugation at 6,000 rpm for 5 min. The supernatant was removed and the pellet was resuspended in RPMI medium (PAN-Biotech GmbH, Germany) supplemented with 10 % fetal bovine serum (FBS) (Gibco Life Technologies, France). The suspension was incubated at 37°C, 5 % CO_2_. An aliquot of culture was collected at 1 and 2 hours. The bacterial load was determined in unheated samples (corresponding to the number of total bacteria, i.e. spores and bacilli) and in heat-treated samples to kill vegetative cells (30 minutes at 65°C; allowing the determination of the number of spores). 100 μL of pure samples or serially diluted samples was plated on BHI agar. At the same time points, an additional volume of culture was filtered through Ultrafree-MC centrifugal Filter Units 0.22 µm (Millipore, France) and stored at -20 °C for ulterior LF and EF expression analysis.

### Infection of mice

Eight- to nine-week-old female C57BL/6J mice (Charles River, France) were maintained in BSL3 facilities. Anesthetized mice with isoflurane were challenged with spores of Bcbva CA into the ear pinna as described previously (27). The inoculum size was verified retrospectively by plating 10-fold serial dilutions on BHI agar plates (4.27 ± 0.30 log10 CFU - mean ± SD, n=6).

At the various stages of the challenge (1 hour, 18 hours and 24 hours), no suffering was induced and no symptom of disease was observed.

### Bacterial enumeration and germination

The presence of vegetative bacilli and spores was determined in the injected ear, blood, spleen, cervical lymph nodes (cLNs) draining the injected ear, liver, lungs and kidneys at 1, 18 and 24 hours post-infection. To assess bacterial germination in these tissues, mice were divided into two groups for each experiment, as previously described [36]. Briefly, for one group, CFU counts were determined directly after organ homogenization in ice-cold distilled water supplemented with 10 % protease inhibitor cocktail (Merck, France), allowing determination of the total bacterial load, i.e. spores and germinated spores. For the other group, organs collected in distilled water supplemented with 10 % protease inhibitor cocktail were heated after collection at 65 °C for 30 minutes and then homogenized. Heat treatment kills vegetative bacteria, allowing determination of the number of spores. Bacterial enumeration was performed by plating pure samples and 10-fold serial dilutions onto BHI agar plates.

For each experiment, six animals were challenged at the same time and randomly allocated to the two groups at the time of sampling. Experiment was repeated two times.

The limits of detection were 40 CFU for lungs and liver, 5 CFU for ear, spleen and kidneys, 2 CFU for cLNs and 10 CFU for blood.

The samples (homogenized ear, cLNs, spleen, blood) were centrifuged at 7,000 rpm for 4 min. The liver and lungs, which were homogenized in larger volumes, were centrifuged at 5,500 rpm at 4°C for 20 min. The plasma and organ samples were then filtered on Ultrafree-MC centrifugal Filter Units 0.22 µm or on syringe filter 0.2 µm for lungs and liver (Merck, France). The filtered samples were stored at -80 and -20 °C for ulterior analysis.

### EF enzymatic activity assay

The adenylyl cyclase EF activity was measured by assessing production of cAMP using a competitive EIA, as previously described, in the plasma, homogenized injected ear, spleen, lungs, cLNs and RMPI-SVF [37,38]. Briefly, the EF adenylyl cyclase activity was first assayed directly in the sample of different tissue (ear, plasma, lungs, spleen and cLNs) and RPMI-SVF medium for *in vitro* experiments, by the enzymatic reaction assay. ATP interference was eliminated and the sensitivity of cAMP detection increased by chemical transformation, followed by the EIA assay. The absorbance was recorded at 405 nm using a spectrophotometer microplate reader iMark (BIO-RAD).

The limit of detection (LOD) was determined as the mean of the negative controls plus three times their standard deviation (SD). Thus, LOD was 0.76 pg/mL in plasma, 1.04 pg/mL in ear, 0.03 pg/mL in lungs, 1.04 pg/mL in cLNs, 0.76 pg/mL in spleen and 0.48 pg/mL in RPMI-SVF. A minimum of 10 negative controls for each matrix was used for the calculation.

### LF enzymatic activity assay

LF ELISA was developed by Nathalie Morel et al. (Laboratoire d’études et de recherches en immunoanalyse, Institut des Sciences du Vivant Frédéric Joliot, CEA, Saclay, France). Briefly, each well of high-binding microtiter plates (Maxisorp Nunc, Dominique Dutscher, France) was coated with 10 μg/mL of a recombinant LF in potassium phosphate buffer 50 mM, at room temperature, overnight. Wells were washed with phosphate potassium buffer containing 0.05 % Tween 20 and blocked with 0.1 % casein in PBS-azide 0.01 %. Plates were conserved at +4°C until their use.

Before their use, wells were washed. Standards were prepared by serial dilution of recombinant LF from 250 to 3.9 pg/mL. Samples (plasma, homogenized ear, cLNs, spleen, lungs and the result of spore germination in RPMI-SVF), negative and positive controls were prepared in adequate dilutions to ascertain to be within the standard curve and incubated in duplicate together with the standards for 35 min at room temperature, under agitation. After washing the wells, biotinylated secondary antibody was used in a concentration of 100 ng/mL for 35 min at room temperature, under agitation. After washing, wells were incubated with poly-horse radish peroxidase (dilution 1:20,000) for 30 min, under agitation. After ultimate washing, plates were developed in the dark with 50 μL of TMB substrate (Thermo Fisher Scientific, France) per well for 30 min and stopped with 50 μL of H_2_SO_4_ (2N). Absorbance was measured at 450 nm (reference wavelength 620nm) using a spectrophotometer microplate reader iMark (BIO-RAD).

The LOD was determined as the mean of the negative controls plus three times their SD. Thus, LOD was 4.2 pg/mL in the plasma, 2.79 pg/mL in the ear, 4.29 pg/mL in the lungs, 2.49 pg/mL in the cLNs, 2.41 pg/mL in the spleen and 1.77 pg/mL in RPMI-SVF medium. A minimum of 12 negative controls for each matrix was used for the calculation.

### Hayluronic acid production and plasma chemistry

Soluble hyaluronic acid (HA) was quantified in the plasma and injected ear using a hyaluronan quantikine ELISA kit, following manufacturer’s instructions (Bio-Techne, France).

Plasma analysis was performed for alanine aminotransferase (ALT; FineTest, Zytomics, France), mouse cardiac Troponin I type 3 (Thermo Fisher Scientific, France) and mouse creatinine (Abedio, Zytomics, France) following manufacturer’s instructions.

### Statistical analysis

Statistical analysis and graphing were performed with GraphPad Prism software (version 4.0; GraphPad Software, San Diego, CA). Statistical significance was determined using Student’s t-test, with a statistical power of 80% at 5% level significance. P < 0.047 was set as the statistically significant threshold.

The lethal dose 50 (LD_50_) value of *G*. *mellonella* larvae infected with Bcbva CA was determined using probit analysis with 95% confidence intervals (nonlinear regression analysis, GraphPad Prism software, version 4.0).

## 3. Results

### 3.1. In vitro production of LF and EF

The secretion of LF and EF by Bcbva CA was quantified *in vitro* after germination and produced amounts were compared to those of *B. anthracis* 9602-*lux* (Figure 1). One hour after culture, for Bcbva CA (6.14 log10 CFU/mL, mean, n=13; Table 1), the mean levels of EF and LF were 32 pg/mL (n=6) and 2.15 x 10^3^ pg/mL (n=4) respectively. For *B. anthracis* 9602-*lux* (6.45 log10 CFU/mL, mean, n=6; Table 1), the mean EF level was 22.8 pg/mL (n=3) and the mean LF level was 2.16 x 10^3^ pg/mL (n=4). There were not significant differences between the two strains. Two hours after culture, there were no longer significant differences. For CA, the mean EF and LF values were 481 pg/mL (n=11) and 1.43 x 10^4^ pg/mL (n=7) respectively, and for 9602-*lux*, the mean values were 526 pg/mL for EF (n=6) and 1.25 x 10^5^ pg/mL for LF (n=4).

**Figure 1.**
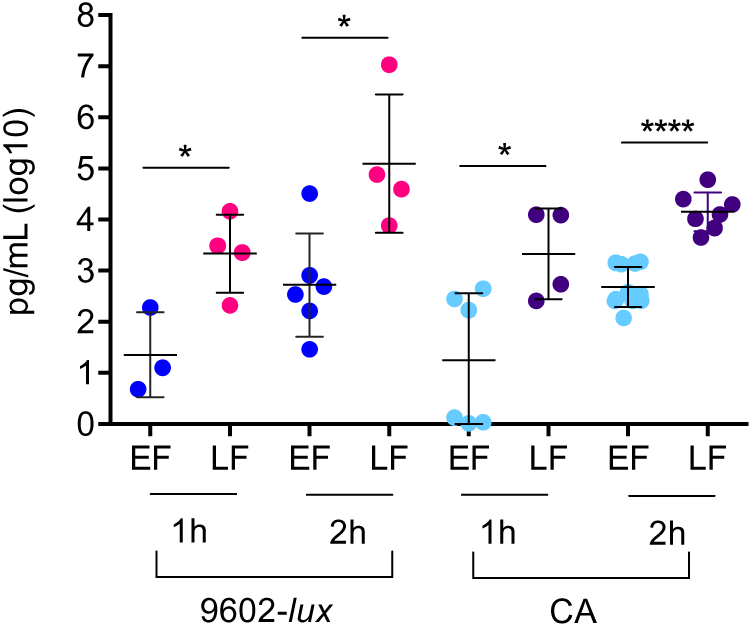
Production of LF and EF *in vitro*. Germination of *B. anthracis* 9602-*lux* and Bcbva CA spores was induced in BHI broth for ten minutes. Bacterial pellets were suspended in RPMI medium supplemented with 10% fetal bovine serum. Bacterial suspension was incubated at 37°C, 5 % CO_2_. At 1 and 2 hours, EF and LF production was quantified as described in the section Materials and methods. Results are expressed as log_10_ pg per mL; each dot represents an individual experiment and the bar represents the mean for each EF and LF quantification at 1 and 2 hours. Each experiment was repeated four to eleven independent times. Only positive values are presented. Asterisks denote statistically significant differences (student t-test); *p < 0.047; **p < 0.01; ***p < 0.001.

**Table 1.** Bacteria level for the strains *B. anthracis* 9602-*lux* and Bcbva CA *in vitro*.

| Time after induction of germination | CFU (log<sub>10</sub>/mL) 9602-*lux* | CFU (log<sub>10</sub>/mL) CA |
| --- | --- | --- |
| 1 hour | 6.45 ± 0.17<br>(6) | 6.14 ± 0.08<br>(13) |
| 2 hours | 7.10 ± 0.18<br>(6) | 6.34 ± 0.09<br>(13) |

Data are expressed as log<sub>10</sub> mean ± SEM, number of experiments (n) in brackets.
Abbreviation: CFU, colony-forming units.

### 3.2. Dose-dependent mortality of Galleria mellonella larvae infected with Bcbva CA

To assess the virulence of spores used in the study, we tested the sensitivity of *Galleria mellonella* larvae model to infection [17]. PBS injection did not induce any mortality (Figure 2). The highest doses tested, 5 x 10^5^ and 2.5 x 10^5^ spores of CA, killed all larvae, within 24 hours. The effect of CA was dose-dependent: with 4.43 x 10^4^ spores injected, the survival rate was 21.7%. It increased to 80.6% with 4.06 x 10^3^ spores injected and to 90% with 4.45 x 10^2^ spores, 10 days post-infection. The LD_50_ was estimated at 2.00 x 10^4^ spores.

**Figure 2.**
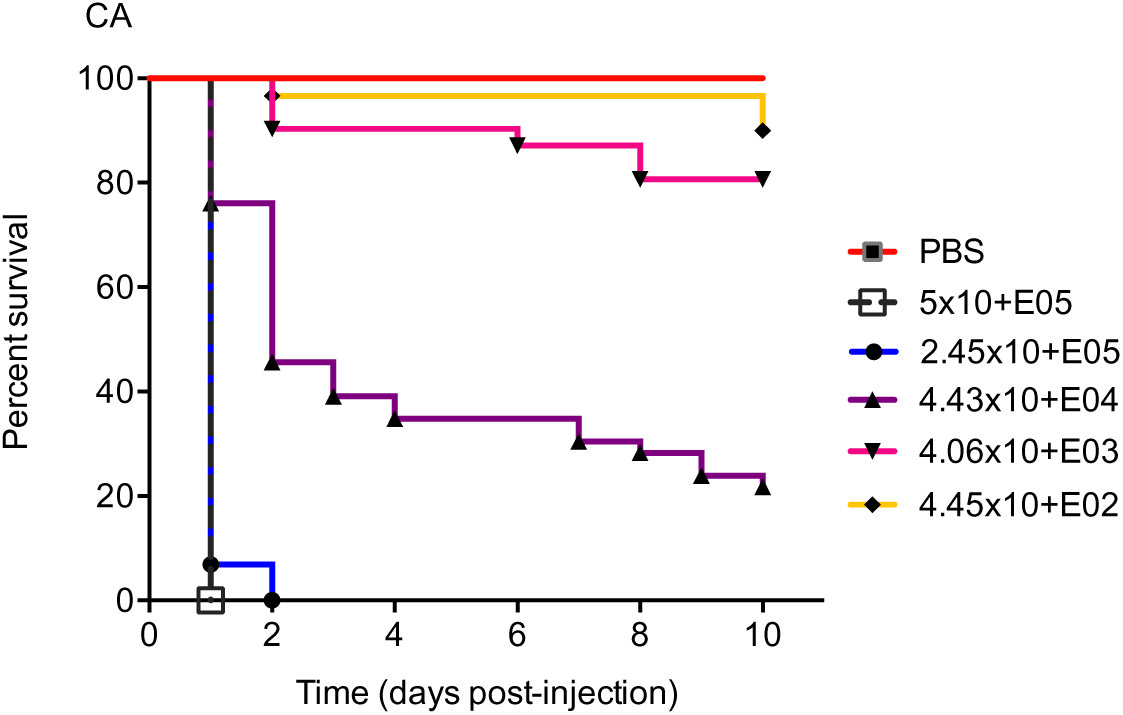
Virulence of the Bcbva strain CA in the *Galleria mellonella* larvae model. Larvae were injected with different doses of spores of CA or with PBS (10 μL) for the group control, as described in the section Materials and Methods. 10 to 12 larvae were assigned to each dose. Each experiment was repeated three independent times. Larvae were monitored over 10 days for survey. Percent survival in time is presented as Kaplan-Meier survival curves.

### 3.3. Rapid bacterial germination and production of LF and EF at the initial site of cutaneous infection with Bcbva CA

To evaluate the dynamics of the toxi-infection associated with Bcbva, immuno-competent mice were infected cutaneously with a fully virulent wild-type Bcbva strain, CA, that produces PDGA and HA capsules, and the toxins LT and ET. We analyzed three times of infection, 1 hour, 18 and 24 hours.

To determine germination, we compared the total bacteria (spores and germinated spores) and spores (heat-resistant bacteria). At each stage of infection, quantification of LF, EF and HA capsule was performed in the homogenized inoculated ear tissue. We observed 79% of germination of CA in the injected ear 1 hour post-infection (Table 2), with 2.56 x 10^3^ total bacteria per ear (NH condition, spores and bacilli; mean, n=6) and 711 spores/ear (H condition, mean, n=6) (Figure 3A). Germination was associated with production of LF, at circa 818 pg/ear (mean, n=6) while EF was not detected as early (Figure 3B, 3C). Bacterial load and germination significantly increased at 18 hours (98%) and 24 hours (100%) (Table 2), with 2.15x10^5^ total bacteria/ear (mean, n=6) and 277 spores (mean, n=6) 24 hours post-infection. LF concentration increased to circa 1.09 x 10^4^ pg/ear at 18 hours and 2.31 x 10^4^ pg/ear (mean, n=6) 24 hours post-infection. EF was detected in all mice at 18 hours, with an average concentration of 762 pg/ear (mean, n=6). Its level hightly increased to 4.43 x 10^3^ pg/ear (mean, n=6) at 24 hours. The mean LF/EF ratio was 14 at 18 hours (n=6) and it decreased to 5 at 24 hours (mean, n=6) (Figure 3D). HA was detected in all mice 18 hours post-inoculation, at a mean level of 2.03 x 10^3^ ng/mL (n=6) (Figure 3E). Its level was significantly higher than that of mice inoculated with PBS but it did not vary between 18 and 24 hours.

**Figure 3.**
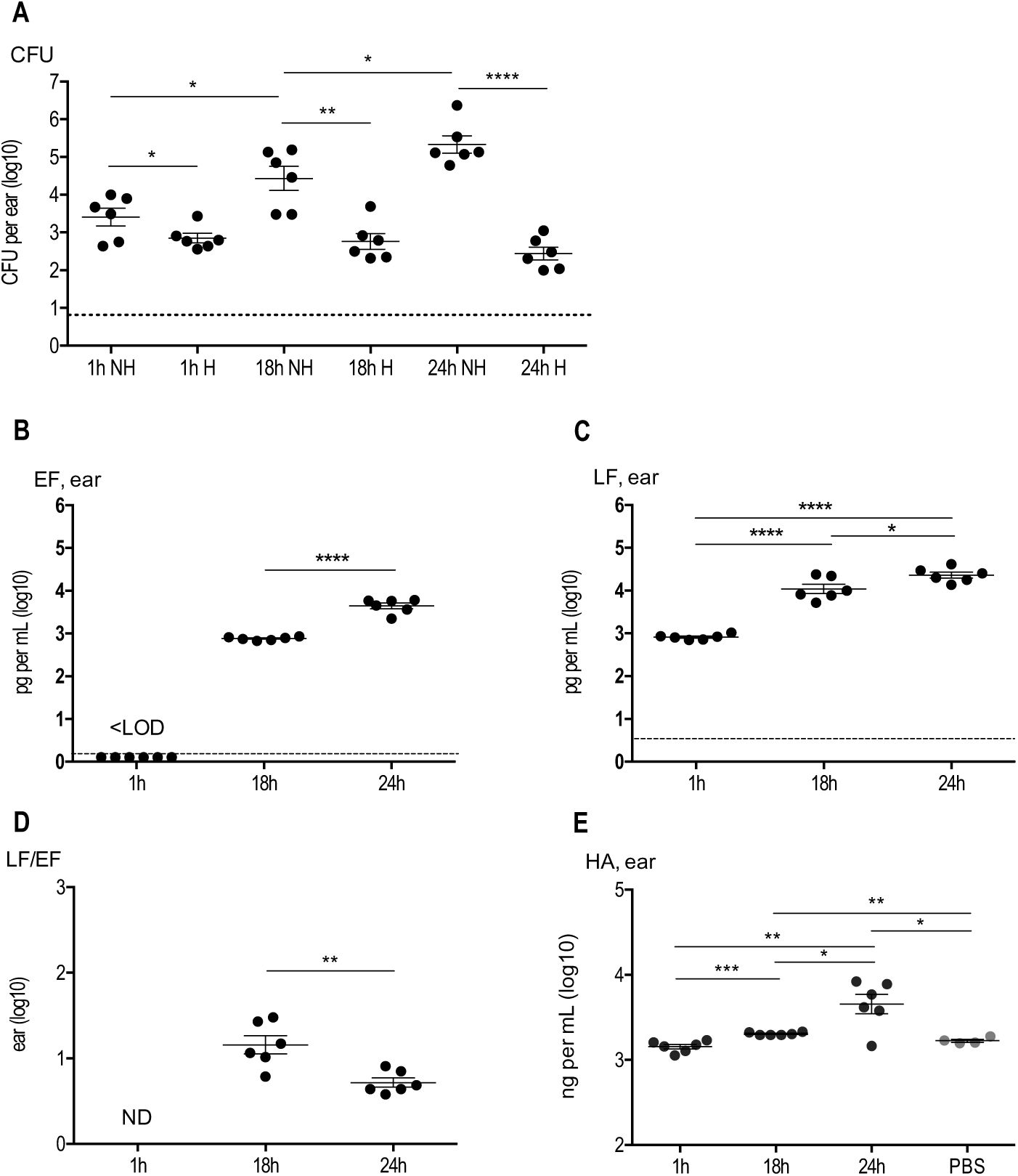
*In vivo* dynamics of EF, LF and HA production at the initial site of infection with Bcbva CA. C57BL/6J mice were infected into the ear pinna with spores of Bcbva CA (4.27 ± 0.30 log10 -mean ± SD, n=6-) as described in the section Material and Methods. Bacterial load (**A**), the levels of edema factor (EF, **B**), lethal factor (LF, **C**) and hyaluronic acid capsule (HA, **E**) were quantified in the homogenized inoculated ear tissue as described in the section Materials and methods, for each time of infection, 1 h, 18 h and 24 h. CFU counting was directly determined after homogenization of organs, to obtain total CFU, i.e. spores and germinated spores (NH for not heated) or was determined after heat treatment and homogenization of organs to get spores (H for heated). The LF/EF ratio (**D**) was calculated from LF and EF values obtained for each individual mouse. Threshold values are represented as dotted lines (0.9 log_10_ for CFU, 0.02 log_10_ pg/mL of homogenized ear and 0.4 log_10_ pg/mL of homogenized ear for EF and LF respectively). Results are expressed as log_10_ pg per mL of homogenized ear for EF and LF and log_10_ ng per mL for HA; each dot represents an individual mouse and the bar represents the mean for each mouse population at each infection stage. Asterisks denote statistically significant differences (student t-test); *p < 0.047; **p < 0.01; ***p < 0.001.

**Table 2.**
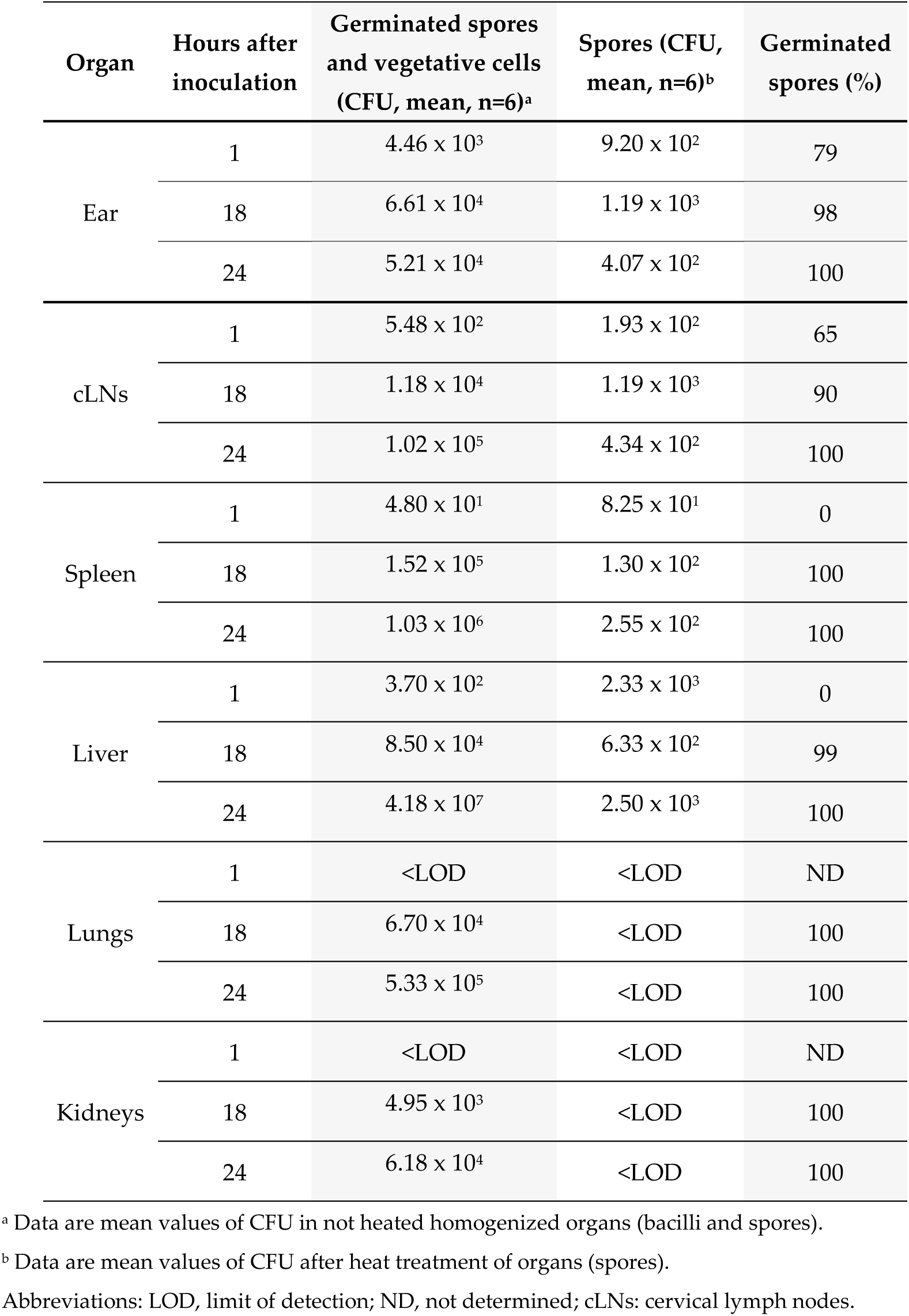
Germination of Bcbva CA in different organs in infected mice.

Germination was fast and important in the injected ear. LF was thus detected very early locally, whereas EF and HA were detected later. LF level was always higher than that of EF, but the ratio LF/EF decreased in time.

### 3.4. Very rapid dissemination of Bcbva CA into lymphoid organs and liver

At the early stage of infection, 1 hour after the challenge of mice, bacteria were detected in the cervical lymph nodes (cLNs) draining the inoculated ear of all mice (Figure 4A). Bacterial loads were heterogeneous between mice, with a minimum value of 2 bacteria detected in the cLNs and a maximum value of 4.33 x 10^3^ bacteria for another mouse. Germination was already significant, with a mean value of 65% (n=6) (Table 1). Bacteria were also detected in the spleen in 83% of inoculated mice, with a low but more homogeneous level of circa 37 CFU (mean, n=5) (Figure 4B). Liver of 67% of mice displayed bacteria, with a higher mean level of 324 bacteria (n=4) (Figure 4C). No germination was highlighted 1 hour post-infection in the spleen and liver.

**Figure 4.**
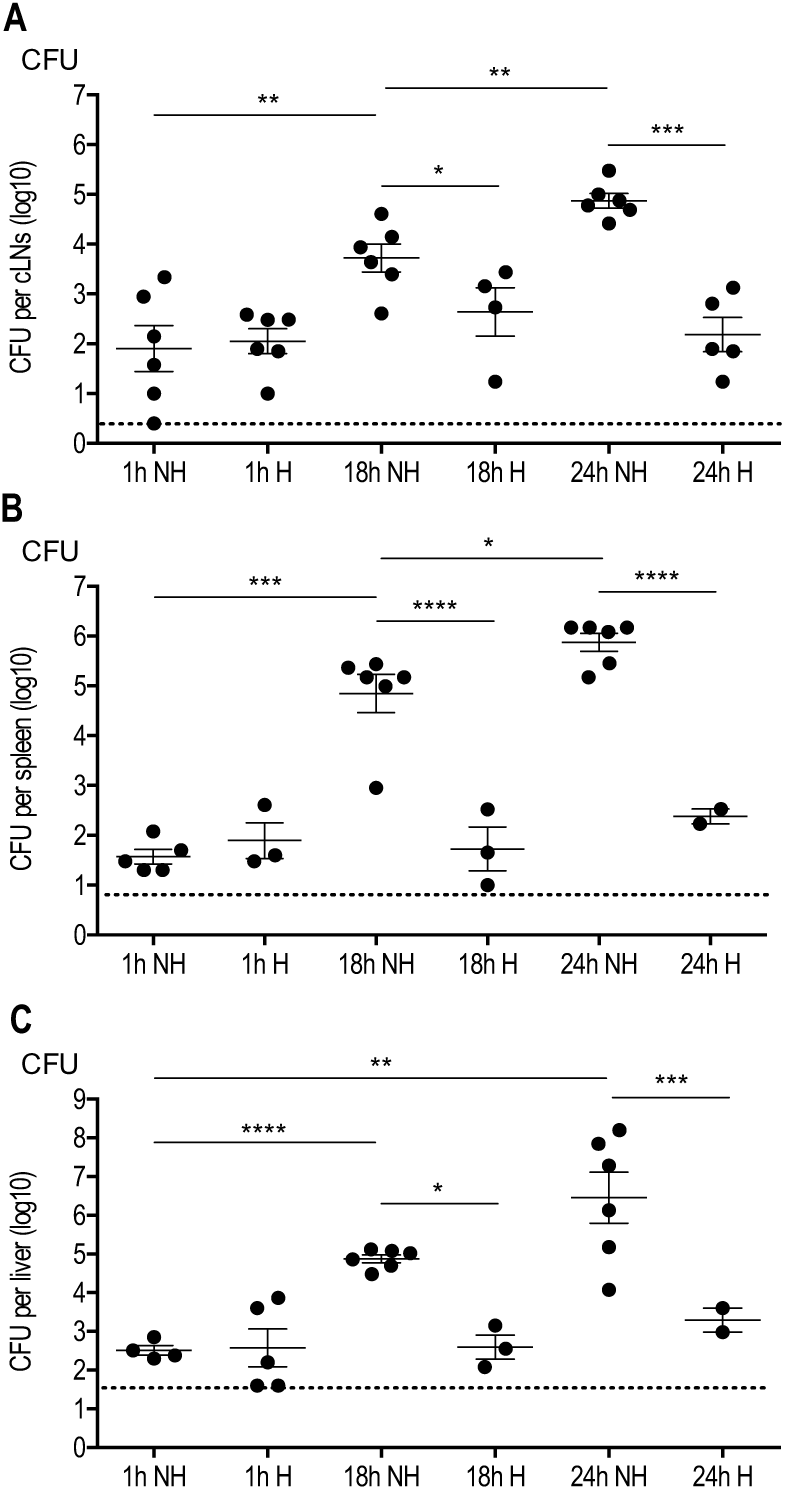
Very rapid dissemination of bacteria into lymphoid organs and liver after cutaneous infection with Bcbva CA. Same experimental conditions as described in Figure 3. For each infected mice, the draining cervical lymph nodes (cLNs, **A**), spleen (**B**) and liver (**C**) were collected. Bacterial CFU was determined in organs without heat treatment (NH) or with heat treatment (H) for each time of infection. Threshold values are represented as dotted lines (0.3 log_10_ for cLNs, 0.7 log_10_ for spleen and 1.6 log_10_ for liver). Results are expressed as log_10_ per total tissue; each dot represents an individual mouse and the bar represents the mean for each mouse population at each infection stage. Asterisks denote statistically significant differences (student t-test); *p < 0.047; **p < 0.01; ***p < 0.001.

Bacterial dissemination in the spleen and liver was observed for all mice 18 hours post-infection. The bacterial load significantly increased between 1 and 24 hours. The level of bacteria reached a mean value of circa 9.4 x 10^4^ CFU in the cLNs, 7.5 x 10^5^ CFU in the spleen and 2.9 x 10^6^ CFU in the liver at 24 hours (n=6 for each organ) (Figure 4B, 4C). Germination increased in the cLNs (90 %, Table 1) and appeared in the spleen (100%, Table 1) and liver (99%, Table 1) 18 hours post-infection, and was 100% in the three organs 24 hours after inoculation.

We highlighted rapid bacterial dissemination in lymphoid organs and liver. Germinated spores were first observed in the cLNs and then, in the spleen and liver in all mice at the final stage of disease.

### 3.5. Early detection of LF in the cLNs and spleen during cutaneous infection with Bcbva CA

One hour after infection, although 65 % of mice displayed bacteria in their cLNs, all mice were positive for LF and EF detection in this compartment (Figure 5). LF level was circa 173 pg/mL and EF level was more than ten times lower, with 13 pg/mL (mean, n=6). Despite the absence of germination in the spleen at this stage of infection, all mice displayed LF in this lymphoid organ (n=6, Table 1) with a high mean value of 2.12 x 10^3^ pg/mL; EF was not detected (n=0/6).

**Figure 5.**
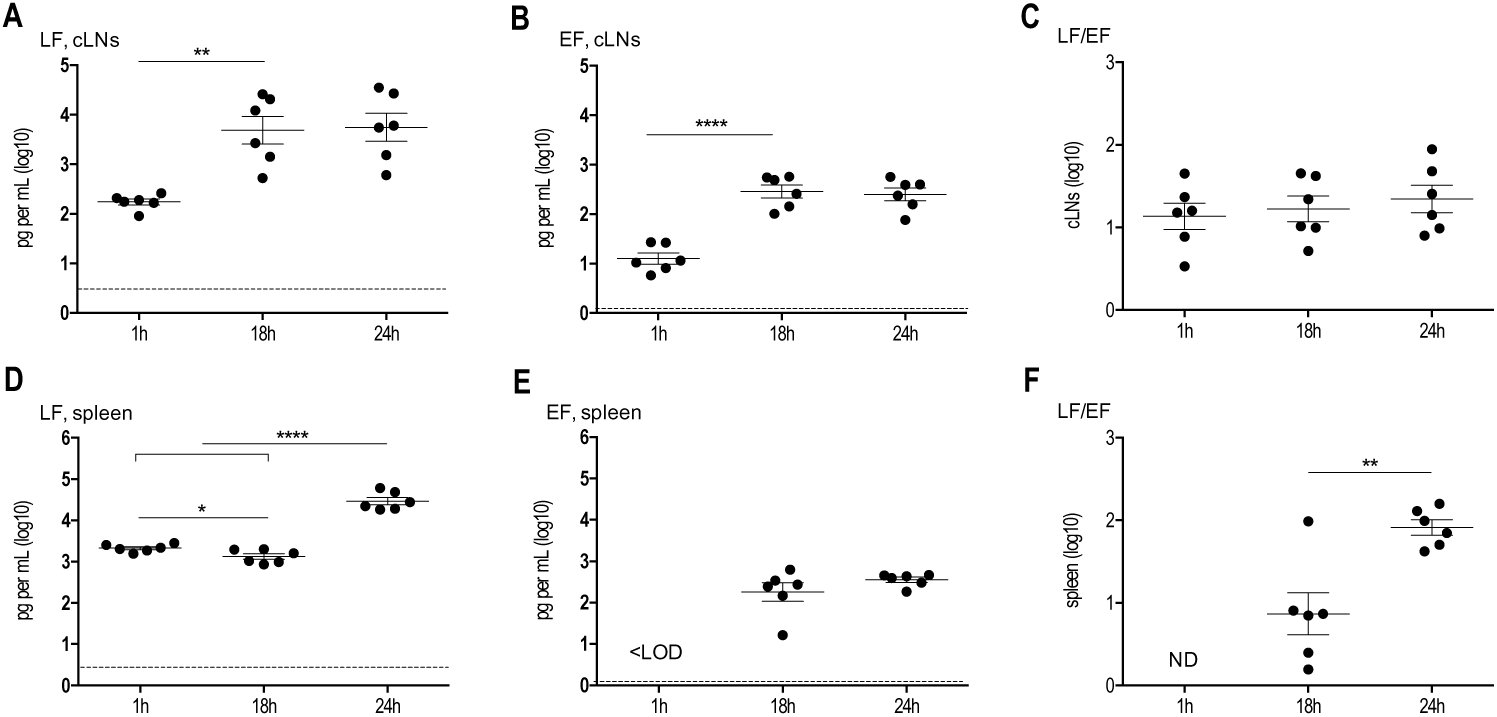
*In vivo* dynamics of EF and LF detection in lymphoid organs during cutaneous infection with Bcbva CA. Same experimental conditions as described in Figures 3 and 4. For each infected mouse, LF (**A, D**) and EF (**B, E**) were quantified in the homogenized cLNs (**A, B**) and liver (**C, D**). The LF/EF ratio (**C, F**) was calculated from the LF and EF values obtained for each individual mouse. Threshold values are represented as dotted lines (0.4 log_10_ pg/mL of homogenized cLNs and 0.38 log_10_ pg/mL of homogenized spleen for LF; 0.02 log_10_ pg/mL of homogenized cLNs and -0.12 log_10_ pg/mL of homogenized spleen for EF). Results are expressed as log_10_ pg per mL of homogenized tissue for EF and LF; each dot represents an individual mouse and the bar represents the mean for each mouse population at each infection stage. Asterisks denote statistically significant differences (student t-test); *p < 0.047; **p < 0.01; ***p < 0.001.

In the cLNs, LF and EF displayed the same dynamic profile during infection, with an increase of their level between 1 and 18 hours, and a plateau between 18 and 24 hours (Figure 5A, 5B). This similar dynamic profile explained the constant value of the ratio LF/EF, with a mean level circa 17 (mean for the 3 times, n=18) (Figure 5C). 18 hours post-inoculation, LF level was 4.8 x 10^3^ pg/mL and EF level was 287 pg/mL (mean, n=6). After 24 hours, LF concentration was 5.6 x 10^3^ pg/mL and EF level was circa 250 pg/mL (mean, n=6) (Figure 5A, 5B).

In the spleen, EF amount did not vary significantly between 18 and 24 hours (180 pg/mL and 356 pg/mL respectively; mean, n=6). The concentration of LF slightly decreased to 1.32 x 10^3^ pg/mL at 18 hours (mean, n=6) and then significantly rose to 2.92 x 10^3^ pg/mL at 24 hours (mean, n=6) (Figure 5D, 5E). Consequently, the LF/EF ratio was high at 24 hours, with a mean value of 82 (Figure 5F).

The presence of LF and EF in these organs early in the infection while germination was not complete or not detected showed a rapid diffusion of the toxin factors far from their site of production.

### 3.6. Early systemic dissemination of bacteria during the course of cutaneous infection

We analyzed the dissemination of total bacteria in the blood circulation and of total bacteria and spores in the lungs and kidneys.

No bacteria were detected in the three compartments 1 hour after infection (Figure 6). 18 hours post-infection, all mice displayed bacteria in the blood (Figure 6A) and interestingly, only bacilli were detected in the lungs and kidneys (Figure 6B, 6C, Table 1), at a mean value circa 1.33 x 10^3^ CFU/mL in the blood (n=12) (Figure 6A), 3.92 x 10^4^ CFU/lungs (Figure 6B) and 3.3 x 10^3^ CFU/kidneys (Figure 6C) (n=6 for each tissue; germination of 100 % in the lungs and kidneys). 24 hours post-infection, only germinated spores were still detected in the lungs and kidneys, with higher mean values of 3.01 x 10^5^ CFU/lungs and 3.32 x 10^4^ CFU/kidneys. In the blood, the bacterial load also significantly increased to a mean value of 1.32 x 10^5^ CFU/mL (n=12).

**Figure 6.**
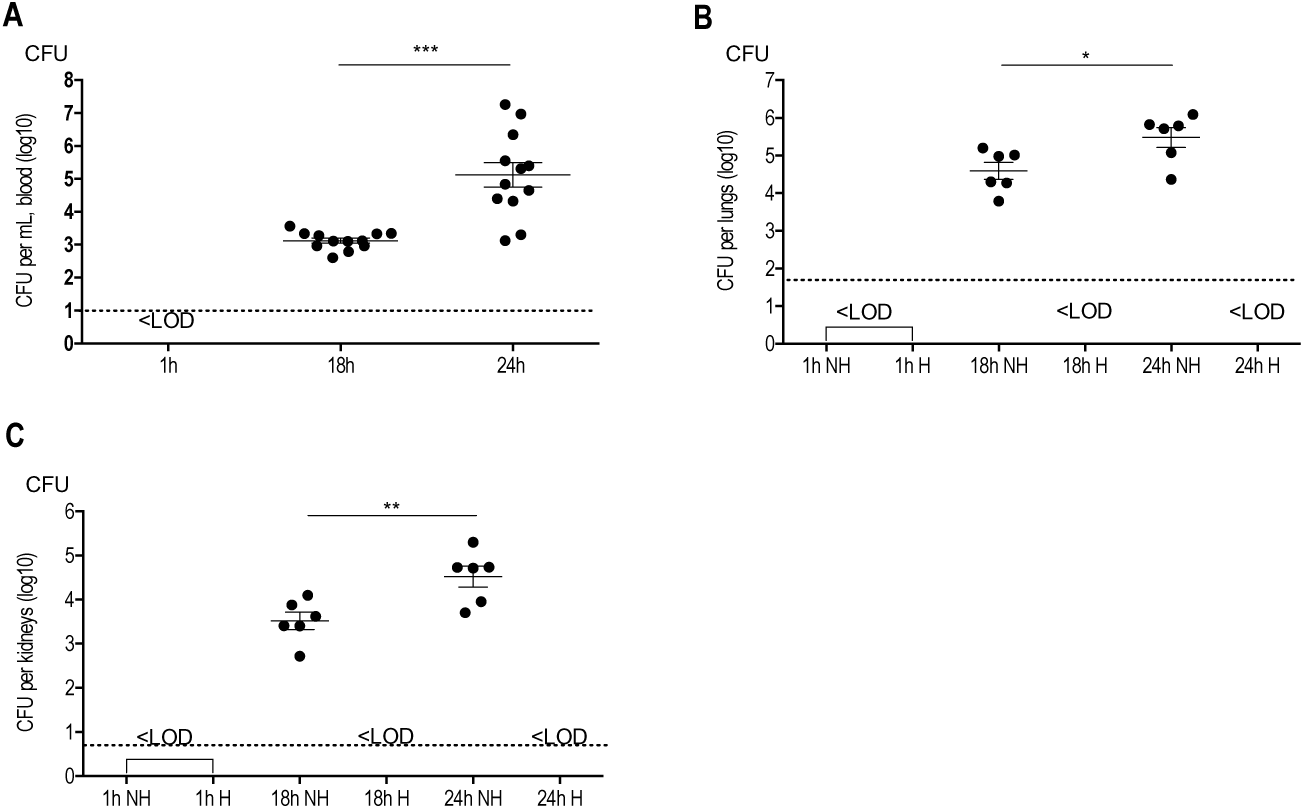
Early systemic bacterial dissemination after cutaneous infection with Bcbva CA. Same experimental conditions as described in Figure 2. For each infected mice, the blood (**A**), lungs (**B**) and kidneys (**C**) were collected. Bacterial load was determined in organs without heat treatment (NH) and with heat treatment (H) for each time of infection. The blood was not heat-treated. Threshold values are represented as dotted lines (1 log_10_ for the blood, 1.6 log_10_ for the lungs and 0.7 log_10_ for the kidneys). Results are expressed as log_10_ per total tissue and as log_10_ per mL for the blood; each dot represents an individual mouse and the bar represents the mean for each mouse population at each infection stage. Asterisks denote statistically significant differences (student t-test); *p < 0.047; **p < 0.01; ***p < 0.001.

The results showed that cutaneous infection with the Bcbva strain CA led to a rapid septicemia of mice.

### 3.7. Systemic diffusion of LF during the course of cutaneous infection

Although no bacteria were detected in the blood samples 1 hour after cutaneous challenge, LF was detected in all mice (n=6 of 6) (Figure 7A), demonstrating a rapid systemic diffusion of LF. EF was not detected as early (n=0 of 6) (Figure 7B). Plasma LF level was 3.94 x 10^2^ pg/mL (mean, n=6) 1 hour after cutaneous infection and it hightly increased 18 and 24 hours after infection to 1.27 x 10^4^ pg/mL (mean, n=6) and 4.32 x 10^5^ pg/mL (mean, n=6) respectively (Figure 7A). All mice showed blood-circulating EF at 18 hours, at a level of 63.2 pg/mL (mean, n=6) (Figure 7B). EF concentration strongly increased at 24 hours with a mean value of 2.51 x 10^3^ pg/mL (n=6). It corresponded to a mean LF/EF ratio of 296 and 172 at 18 and 24 hours (Figure 7C).

**Figure 7.**
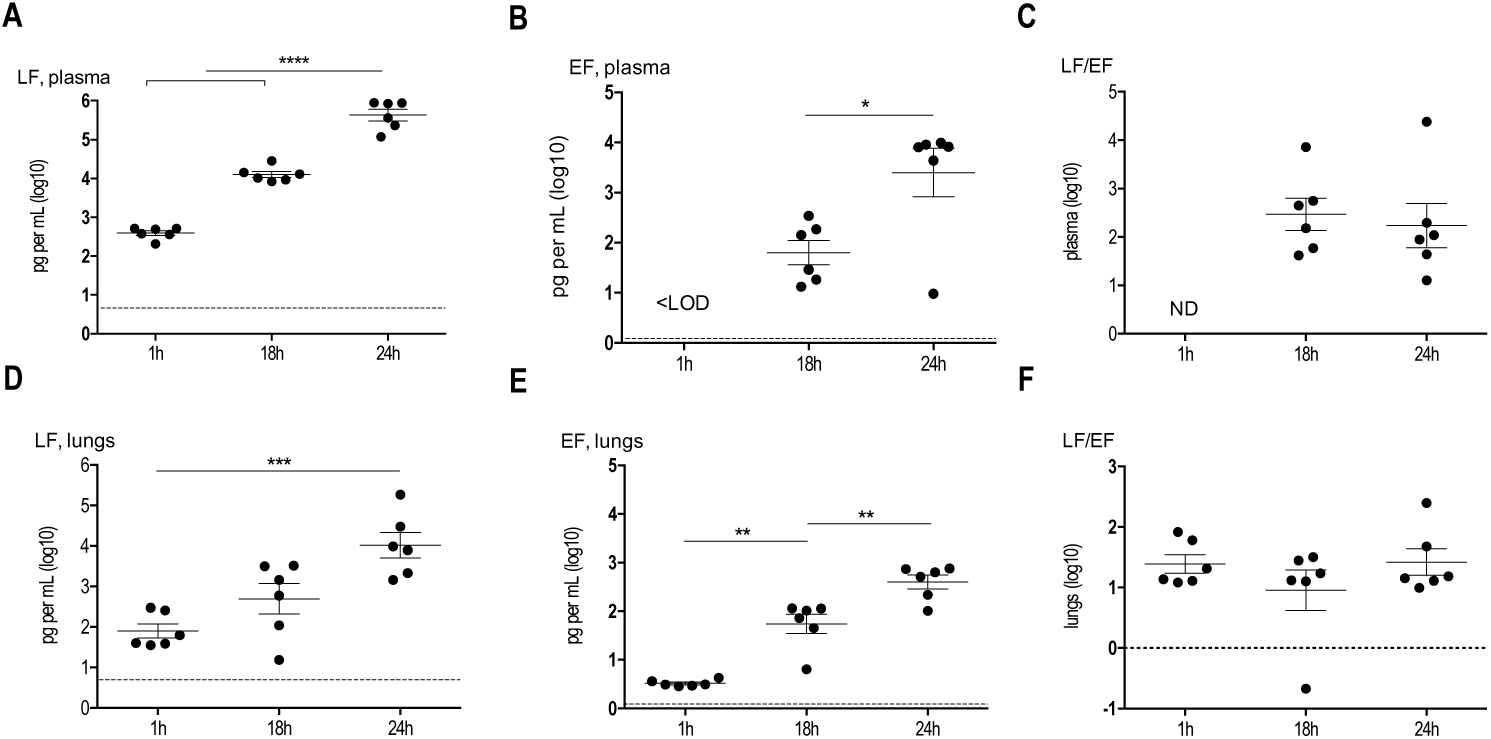
*In vivo* dynamics of LF and EF detection in the plasma and lungs during cutaneous infection with Bcbva CA. Same experimental conditions as described in Figures 3 and 4. For each infected mouse, LF (**A, D**) and EF (**B, E**) were quantified in the plasma (**A, B**) and homogenized lungs (**D, E**). The LF/EF ratio (**C, F**) was calculated from the LF and EF values obtained for each individual mouse. Threshold values are represented as dotted lines (0.62 log_10_ pg/mL in the plasma and 0.63 log_10_ pg/mL of homogenized lungs for LF; -0.12 log_10_ in the plasma and -1.47 log_10_ pg/mL of homogenized lungs for EF). Results are expressed as log_10_ pg per mL of homogenized tissue for EF and LF; each dot represents an individual mouse and the bar represents the mean for each mouse population at each infection stage. Asterisks denote statistically significant differences (student t-test); *p < 0.047; **p < 0.01; ***p < 0.001.

Surprisingly, all mice presented LF and EF in the lungs, 1 hour after infection, while no bacteria were detected (Figure 7D, 7E). As in the blood circulation, it demonstrated a rapid diffusion of the two factors in a part of the respiratory tract. The mean LF concentration was 80 pg/mL (n=6) and EF level was 24 times lower, with 3.3 pg/mL (mean, n=6). Concentration of both factors increased between 1 and 24 hours, with LF level circa 1.04 x 10^4^ pg/mL (mean, n=6) and EF value circa 3.96 x 10^2^ pg/mL (mean, n=6) (Figure 7D, 7E). The LF/EF ratio did not vary significantly at 1, 18 and 24 hours, with a range value from 24 to 26 (Figure 7F).

As observed in lymphoid organs, the systemic presence of LF and EF in the plasma and lungs early in the infection in absence of bacteria highlighted a rapid diffusion of the toxin factors far from their site of production.

### 3.8. Cardiac and renal lesions induced during the course of cutaneous infection

No gross pathology was observed during the collection of the various tissues but to assess more precisely the effect of Bcbva CA infection, we quantified different biomarkers. After infection, no increase of alanine aminotransferase (ALT), a marker of hepatic dysfunction, was observed compared to mice challenged with PBS (Figure 8A). When we analyzed the cardiac troponin (Figure 8B), a significant rise of its level was induced 1 hour and 18 hours after infection, with a mean value of 73 and 75 pg/mL respectively (n=6 for each time), compared to PBS condition (56 pg/mL, mean; n=6). Its concentration highly increased over time, with a mean level circa 205 pg/mL (n=6) at 24 hours post-infection, indicating cardiac damage in the majority of mice, whereas the mean levels in the PBS condition were unchanged between 1 hour and 48 hours (53 pg/mL, mean; n=4). The mean level of creatinine in the plasma of infected mice was higher at 18 hours (4.18 μg/mL; n=5) and 24 hours (5.92 μg/mL; n=5) than that of control mice (1.48 μg/mL for both time points; n=7), suggesting renal impairment.

**Figure 8.**
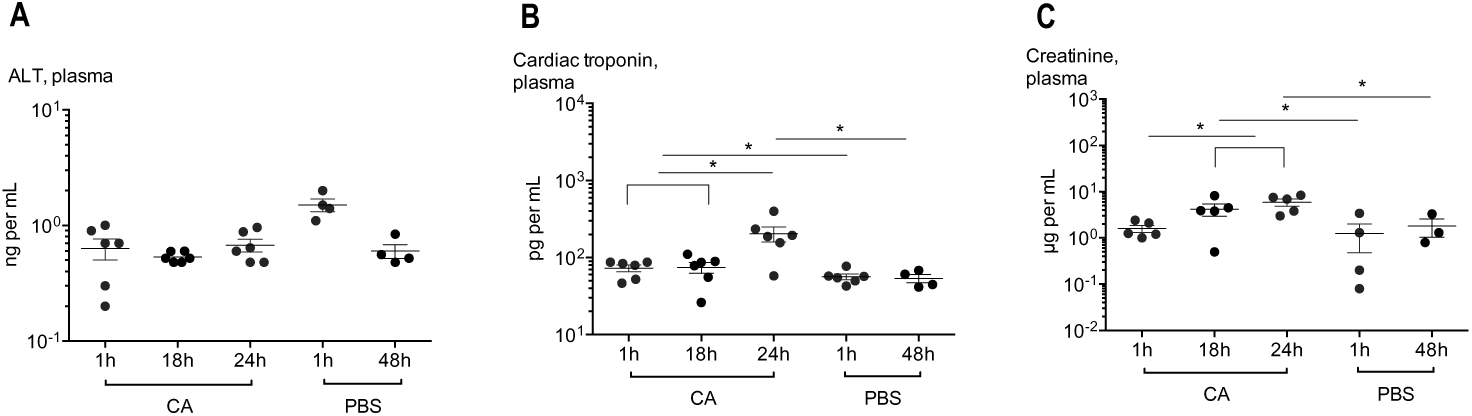
Hepatic, cardiac and renal biomarker levels in the plasma during cutaneous infection with Bcbva CA. Same experimental conditions as described in Figures 3 and 4. For each infected mouse and mice challenged with PBS, alanine aminotransferase (ALT, **A**), cardiac troponin (**B**) and mouse creatinine (**C**) were quantified in the plasma following manufacturer’s instructions. Results are expressed as ng/mL for ALT, pg per mL for cardiac troponin and μg/mL for creatinine; each dot represents an individual mouse and the bar represents the mean for each mouse population at each infection stage. Asterisks denote statistically significant differences (student t-test); *p < 0.047; **p < 0.01; ***p < 0.001.

## 4. Discussion

Few studies have characterized the course of disease associated with Bcbva strains and most have focused on the inhalational route of infection [16,17]. To our knowledge, our study is the first to decipher the toxi-infection associated with the Bcbva strain CA isolated in Cameroon, using an immuno-competent murine model of cutaneous disease. Here, we highlight both similarities and differences with *B. anthracis* cutaneous infection, explaining an equivalent virulence [15].

To reduce the use of vertebrate animals, we first verified the virulence of spores of Bcbva CA on *G. mellonella* larvae. Larvae have already been used for *B. anthracis* and *B. cereus* [39,40] and more recently for Bcbva [17]. As observed for the Bcbva strain CI isolated in Côte d’Ivoire, CA induces a dose-dependent mortality, with a rapid loss of virulence between the dose 4.43 x 10^4^ and 4.06 x 10^3^ spores. We determined a LD_50_ for CA in *G. mellonella* ≈ 22-fold greater than that of Bcbva CI, which presented a LD_50_ of 9.21 x 10^2^ spores [17]. This result differs from the data previously obtained in mice and guinea pigs after subcutaneous injection, where both strains presented a similar LD_50_ [15]. In the study of Jiranantasak, injection of PBS induced a significant mortality, which we did not provoke under our study conditions. Therefore, the level of virulence of CI in *G. mellonella* may be overestimated and it could be closer to what we observe with CA.

In murine models of cutaneous infection, the initial site of infection is a critical point for the future of the disease [41,42]. Spores of *B. anthracis* rapidly germinate at the initial inoculation site [43], as early as 15 minutes after their injection [36] or 1-3 hours after bacterial application onto damaged skin [44]. Germination of spores is associated with the expression of the toxin genes [45] and production of anthrax toxin components [46]. Thereby, LF and EF were produced at an early time point of cutaneous anthrax infection [37]. In the ear inoculated with spores of *B. anthracis* 9602-*lux*, 29% and 38% of mice displayed LF and EF respectively, both factors diffusing rapidly into the blood, despite the absence of circulating bacteria at the early stage of infection [37]. Partly in agreement with these studies, our results highlight Bcbva germination within 1 hour in the injected ear and a local production of LF for all mice, while EF was not detected. Here, we used an ELISA assay to quantify LF, with a limit of detection lower than that the LOD determined for the LC-MS/MS method used in our previous study. It could explain the difference of percentage of mice positive for LF between the two studies. However, the average LF level produced by CA is almost 260-fold lower than that of LF produced by *B. anthracis* in the inoculated ear, at the early phase of disease [37]. *In vitro* study had demonstrated that Bcbva tended to produce less PA than *B. anthracis* Ames [17]. It could have been extrapolated to LF and EF but here, we obtained different results in *in vitro* experiments for both factors. For an equivalent bacterial load, we quantified a similar production of LF and EF for Bcbva compared to *B. anthracis* 9602-*lux*, 1 and 2 hours after culture. Therefore, the ability to produce these factors is not an explanation for the differences observed *in vivo* within 1 hour of infection. We hypothetize that a part of LF produced locally by Bcbva rapidly disseminates into distal compartments. Indeed, 1 hour post-inoculation, LF was detected in the spleen of all mice, at high levels, while there was not germination, and in the lungs and blood despite the absence of bacteria in these compartments. Concerning EF, it is produced at lower levels than LF and is often detected at the same time as, or later than LF in animal models of *B. anthracis* infection [37,38,38,47,48]. Thus, in the ear inoculated with CA, EF may be produced later than LF, or in quantities undetectable with our method. Or, as LF, it may rapidly disseminate into distal organs, as the lungs where it was detected in absence of bacteria and as the cLNs, where it was present. In the ear, we also observe a different profile for LF and EF kinetics compared to that with 9602-*lux*. LF and EF levels increase along infection with Bcbva CA, while LF and EF concentrations rapidly decreased or increased respectively, to reach a plateau, with *B. anthracis* 9602-*lux* [37]. The profiles observed for LF and EF are probably associated with the patterns of bacterial load: a continuous increase of bacterial burden for CA whereas *B. anthracis* presented a short increase at the early phase of disease, followed by a plateau until the final septicemic stage [37]. Nevertheless, early production of toxins are crucial for the colonization of the portal of entry and bacterial dissemination [41,49]. Despite lower levels of LF and EF produced *in vivo* by CA, their early presence presumably allows colonization of the ear and bacterial dissemination [41].

In the study of Ferris, soluble HA capsule was detected in the serum of rabbits at the final stage of inhalational disease, when bacteria were detected in the blood [16]. We obtained a similar result with detection of soluble HA capsule associated with the infection in the plasma 24 hours post-infection (Figure A1). The results obtained in the inoculated ear appeared interesting. We detect soluble hyaluronic acid 18 hours post-infection, with no variation of its concentration between 18 hours and 24 hours. The HA capsule seemed less important in the virulence than the PDGA capsule during subcutaneous infection with CA [15]. The expression of HA not allowed systemic spreading of bacteria but promoted their local multiplication, perhaps by helping them to evade the immune system. Indeed, HA is a key virulent factor expressed in *Streptococcus pyogenes*. Previous studies showed that HA capsule prevented opsono-phagocytic killing of this bacteria by neutrophils and human blood leukocytes [50–52], its uptake by keratinocytes, promoting invasive soft tissue infection [53]. HA may also avoid or limit the transcytosis of bacteria by epithelial cells and fibroblasts observed *in vitro* and *in vivo* for *B. anthracis* [54–56]. HA is also certainly a contributing factor to immune evasion for Bcbva, in synergy with the toxins ET and LT [15,57,58]. Analysis of cytokine response to Bcbva infection and the use of a Bcbva mutant lacking HA expression will respond to many questions. Notably, it may explain the exponential and uncontrolled increase of bacterial burden in the injected ear, with over 80-fold more bacteria in 24 hours. Alternatively, this could simply be explained by the inoculum dose. Indeed, in the study with *B. anthracis* 9602-*lux* in which bacterial growth appeared to be controlled in the injected ear, with only a 20-fold increase in bacterial burden between the early and terminal stages, more than 100-fold fewer bacteria had been injected [37].

In the Jailbreak model, proposed by Weiner and Glosmki [59], toxins and proteases produced at the site of infection allow vegetative bacilli spreading into the regional lymph nodes and then into the blood circulation. Thus, during cutaneous infection, *B. anthracis* bacteria disseminate to the cervical draining lymph nodes (cLNs), the spleen, then the lungs and blood, and finally induce septicemia [36,37,42,43,49]. As *B. anthracis*, we provide evidence that infection with CA rapidly and simultaneously progresses to lymphoid organs and liver, and then, to systemic compartments, while continuing to multiply exponentially at the inoculation site. One hour after infection, approximately 6% of injected bacteria arrived in the cLNs. This percentage has already been observed in a study of Weiner, with 3-4% of injected spores of *B. anthracis* reaching the lymph nodes draining the inoculated ear [42]. However, the proportion of germinated spores of CA in the cLNs 1 hour post-infection is significant. It differs from other studies, where only spores were detected, which suggested a balance between influx of spores and destruction of bacilli [36,42]. Several mechanisms may be involved to explain the presence of germinated spores CA. As already mentionned, the HA capsule, in synergy with EF and LF detected very early in the cLNs, could play a protective role, contributing to avoid the destruction of bacilli by the immune system, when they spread from the ear to cLNs or when germination occurs in the cLNs, and thus, allowing their early presence in this compartment. The immune response in the cLNs deserves investigation to complete the characterization of the infectious process. The HA capsule may also contribute to promote rapid lymphatic dissemination of bacteria, through the interaction with lymphatic endothelial receptor, as demonstrated for encapsulated group A Streptococci [60]. As the proportion of bacteria reaching the cLNs is low, the role of HA in bacterial dissemination did not appear to be essential [15]. Moreover, Bcbva CA strains are motile bacteria, contrary to *B. anthracis* [6]. Motily can contribute to the outbreak of the infection, all mice inoculated with CA developing septicemia within 18 hours.

As observed for *B. anthracis*, CA spores are concurrently detected in the spleen and liver, at low numbers, 1 hour after inoculation [36]. Germination observed in the liver and spleen 18 and 24 hours after infection is associated with a significant increase in the number of bacteria, the spore population remaining stable along the course of disease, as for *B. anthracis* [36]. However, there is no evidence of hepatic dysfunction while anthrax can induce injury of the liver [61–64]. This aspect can be studied in greater detail, by measuring other markers of hepatic function. Mice were in septicemia, in less than a day, displaying increasing bacteremia and only bacilli in lungs and kidneys at 18 hours and 24 hours. These results suggest that CA spores remain trapped in the lymphoid organs, and that only bacilli circulate *via* the bloodstream, to effectively colonize the lungs and kidneys. Associated with the high level of bacteremia and the presence of germinated spores in the kidneys, we observe early cardiac and renal dysfunction, as previously highlighted with *B. anthracis*, with an increase of cardiac troponin and biomarker of renal injury [61,65–71]. It will be interesting to determine the contribution of the different factors of Bcbva CA in these organ failures.

Alongside uncontrolled bacterial multiplication and spreading, our results outline the toxemic part of infection with Bcbva, especially with a rapid production and diffusion of LF. This factor displays a systemic diffusion within 1 hour, in all analyzed tissues while EF is detected in the cLNs and lungs, at lower levels than LF. These results strongly suggest an ability to dissminate far from their site of production. Between 1 hour and 24 hours, the presence of LF and EF in the cLNs and spleen appears more controlled than in the blood and lungs, where their levels exponentially arise. The levels of LF remain higher than the levels of EF, as previously determined for *B. anthracis* [37,38,47,48], making this factor a potential marker for monitoring infections associated with atypical *B. cereus* strains [72]. Early presence of these factors in numerous tissues, particularly LF, their capacity to disseminate and their increasing levels certainly contribute to the fulminant development of the infection and to multi-organ failures.

To conclude, we observe a fast progression of toxi-infection and development of septicemia for all animals, without apparent clinical signs of illness. The rapid evolution of the disease with early cardiac damage and renal impairment may lead to the sudden deaths of wild animals previously observed [2,6,73] and highlights the risk of Bcbva cutaneous infection in humans.

## Author Contributions

Conceptualization, C.R.; methodology, C.R.; formal analysis, C.R. and A.G.; investigation, C.R. and A.G.; resources, C.R.; data curation, C.R.; writing—original draft preparation, C.R.; writing—review and editing, C.R.; project administration, C.R.; funding acquisition, C.R. All authors have read and agreed to the published version of the manuscript.

## Institutional Review Board Statement

The animal study protocol was approved by the SSA animal ethics committee according to applicable French legislation (Directive 2010/63/UE, edict 2013-118, project number 511985, July 9 2024).

## Data Availability Statement

The data presented in this study are available in this article.

## Acknowledgements

We thank Robert Koch-Institute, and particularly Silke Klee, who provided us the Bcbva strain CA (Centre for Biological Threats and Special Pathogens, ZBS 2: Highly Pathogenic Microorganisms, Berlin, Germany). We would like to convey our thanks to Nathalie Morel (Laboratoire d’études et de recherches en immunoanalyse, Institut des Sciences du Vivant Frédéric Joliot, CEA, Saclay, France), for sharing the sensitive LF assay technique. We would like to express our thanks to Christina Nielsen-Leroux and Christophe Buisson for responding to our requests for *Galleria mellonella* (INRAE, Jouy-en-Josas, France). Finally, we thank Clarisse Vigne for the technical support over the weekend (Unité Interaction Hôte-Pathogènes, Institut de Recherche Biomédicale des Armées, Brétigny-sur-Orge, France).

## Abbreviations

The following abbreviations are used in this manuscript:

Bcbva: Bacillus cereus biovar anthracis
PDGA: Poly-γ-D-glutamic acid
EF: Edema factor
LF: Lethal factor
PA: Protective antigen
CFU: Colony forming unit
cLNs: Cervival lymph nodes
LOD: Limit of detection
HA: Hyaluronic acid
LD_50_: Lethal dose 50
h: Hour
min: Minutes
NH: Not heated
H: Heat treatment
ND: Not determined

## Appendix A

**Figure A.1.**
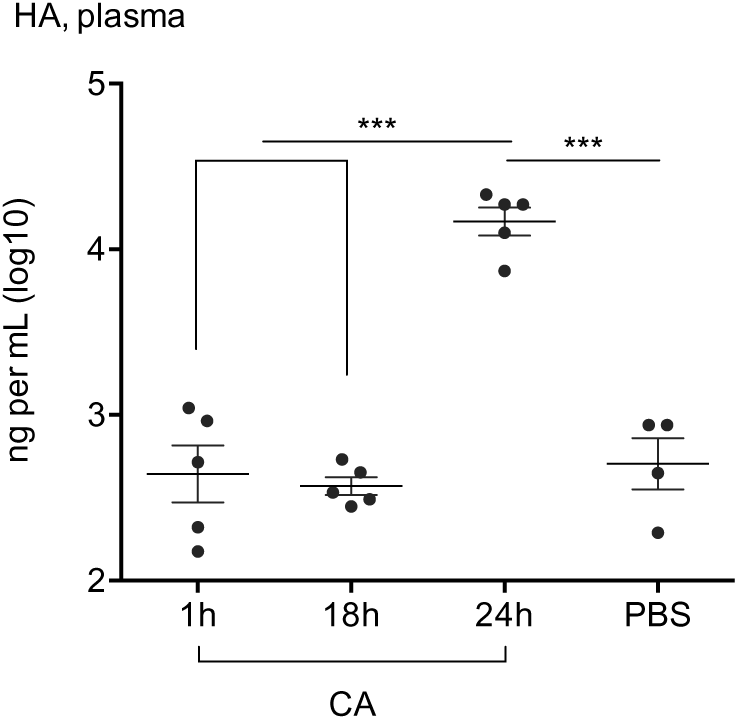
*In vivo* dynamics of HA production in the plasma during cutaneous infection with Bcbva CA. Hyaluronic acid capsule (HA) was quantified in the plasma as described in the Materials and methods section, for each time of infection, 1 h, 18 h and 24 h and for the PBS condition. Results are expressed as log_10_ ng per mL for HA; each dot represents an individual mouse and the bar represents the mean for each mouse population at each infection stage. One color represents one experiment, to see a potential cage effect. Asterisks denote statistically significant differences (student t-test); ***p < 0.001.

## References

1. Jensen, G.B.; Hansen, B.M.; Eilenberg, J.; Mahillon, J. The Hidden Lifestyles of *Bacillus Cereus* and Relatives. Environ Microbiol 2003, 5, 631–640, doi:10.1046/j.1462-2920.2003.00461.x.

2. Leendertz, F.H.; Ellerbrok, H.; Boesch, C.; Couacy-Hymann, E.; Mätz-Rensing, K.; Hakenbeck, R.; Bergmann, C.; Abaza, P.; Junglen, S.; Moebius, Y.;, et al. Anthrax Kills Wild Chimpanzees in a Tropical Rainforest. Nature 2004, 430, 451–452, doi:10.1038/nature02722.

3. Leendertz, F.H.; Lankester, F.; Guislain, P.; Néel, C.; Drori, O.; Dupain, J.; Speede, S.; Reed, P.; Wolfe, N.; Loul, S.;, et al. Anthrax in Western and Central African Great Apes. Am J Primatol 2006, 68, 928–933, doi:10.1002/ajp.20298.

4. Leendertz, F.H.; Yumlu, S.; Pauli, G.; Boesch, C.; Couacy-Hymann, E.; Vigilant, L.; Junglen, S.; Schenk, S.; Ellerbrok, H. A New *Bacillus Anthracis* Found in Wild Chimpanzees and a Gorilla from West and Central Africa. PLoS Pathog 2006, 2, e8, doi:10.1371/journal.ppat.0020008.

5. Klee, S.R.; Ozel, M.; Appel, B.; Boesch, C.; Ellerbrok, H.; Jacob, D.; Holland, G.; Leendertz, F.H.; Pauli, G.; Grunow, R.;, et al. Characterization of *Bacillus Anthracis-*like Bacteria Isolated from Wild Great Apes from Cote d’Ivoire and Cameroon. J Bacteriol 2006, 188, 5333–5344, doi:10.1128/JB.00303-06.

6. Antonation, K.S.; Grützmacher, K.; Dupke, S.; Mabon, P.; Zimmermann, F.; Lankester, F.; Peller, T.; Feistner, A.; Todd, A.; Herbinger, I.;, et al. *Bacillus Cereus* Biovar *Anthracis* Causing Anthrax in Sub-Saharan Africa-Chromosomal Monophyly and Broad Geographic Distribution. PLoS Negl Trop Dis 2016, 10, e0004923, doi:10.1371/journal.pntd.0004923.

7. Zimmermann, F.; Köhler, S.M.; Nowak, K.; Dupke, S.; Barduhn, A.; Düx, A.; Lang, A.; De Nys, H.M.; Gogarten, J.F.; Grunow, R.;, et al. Low Antibody Prevalence against *Bacillus Cereus* Biovar *Anthracis* in Taï National Park, Côte d’Ivoire, Indicates High Rate of Lethal Infections in Wildlife. PLoS Negl Trop Dis 2017, 11, e0005960, doi:10.1371/journal.pntd.0005960.

8. Hoffmann, C.; Zimmermann, F.; Biek, R.; Kuehl, H.; Nowak, K.; Mundry, R.; Agbor, A.; Angedakin, S.; Arandjelovic, M.; Blankenburg, A.;, et al. Persistent Anthrax as a Major Driver of Wildlife Mortality in a Tropical Rainforest. Nature 2017, 548, 82–86, doi:10.1038/nature23309.

9. Norris, M.H.; Zincke, D.; Daegling, D.J.; Krigbaum, J.; McGraw, W.S.; Kirpich, A.; Hadfield, T.L.; Blackburn, J.K. Genomic and Phylogenetic Analysis of *Bacillus Cereus* Biovar *Anthracis* Isolated from Archival Bone Samples Reveals Earlier Natural History of the Pathogen. Pathogens 2023, 12, 1065, doi:10.3390/pathogens12081065.

10. Dupke, S.; Barduhn, A.; Franz, T.; Leendertz, F.H.; Couacy-Hymann, E.; Grunow, R.; Klee, S.R. Analysis of a Newly Discovered Antigen of *Bacillus Cereus* Biovar *Anthracis* for Its Suitability in Specific Serological Antibody Testing. J Appl Microbiol 2019, 126, 311–323, doi:10.1111/jam.14114.

11. Dupke, S.; Schubert, G.; Beudjé, F.; Barduhn, A.; Pauly, M.; Couacy-Hymann, E.; Grunow, R.; Akoua-Koffi, C.; Leendertz, F.H.; Klee, S.R. Serological Evidence for Human Exposure to *Bacillus Cereus* Biovar *Anthracis* in the Villages around Taï National Park, Côte d’Ivoire. PLoS Negl Trop Dis 2020, 14, e0008292, doi:10.1371/journal.pntd.0008292.

12. Kolstø, A.-B.; Tourasse, N.J.; Økstad, O.A. What Sets *Bacillus Anthracis* Apart from Other Bacillus Species? Annu Rev Microbiol 2009, 63, 451–476, doi:10.1146/annurev.micro.091208.073255.

13. Klee, S.R.; Brzuszkiewicz, E.B.; Nattermann, H.; Brüggemann, H.; Dupke, S.; Wollherr, A.; Franz, T.; Pauli, G.; Appel, B.; Liebl, W.;, et al. The Genome of a Bacillus Isolate Causing Anthrax in Chimpanzees Combines Chromosomal Properties of *B*. Cereus with B. Anthracis Virulence Plasmids. PLoS One 2010, 5, e10986, doi:10.1371/journal.pone.0010986.

14. Baldwin, V.M. You Can’t *B*. *Cereus* - A Review of Bacillus Cereus Strains That Cause Anthrax-Like Disease. Front Microbiol 2020, 11, 1731, doi:10.3389/fmicb.2020.01731.

15. Brézillon, C.; Haustant, M.; Dupke, S.; Corre, J.-P.; Lander, A.; Franz, T.; Monot, M.; Couture-Tosi, E.; Jouvion, G.; Leendertz, F.H.;, et al. Capsules, Toxins and AtxA as Virulence Factors of Emerging *Bacillus Cereus* Biovar *Anthracis*. PLoS Negl Trop Dis 2015, 9, e0003455, doi:10.1371/journal.pntd.0003455.

16. Ferris, A.M.; Dawson, D.G.; Eyler, A.B.; Yeager, J.J.; Bohannon, J.K.; Boydston, J.A.; Krause, M.L.; Balzli, C.L.; Wahl, V.; Jenkins, T.D.;, et al. *Bacillus Cereus* Biovar *Anthracis* Causes Inhalational Anthrax-like Disease in Rabbits That Is Treatable with Medical Countermeasures. PLoS Negl Trop Dis 2025, 19, e0012973, doi:10.1371/journal.pntd.0012973.

17. Jiranantasak, T.; Bluhm, A.P.; Chabot, D.J.; Friedlander, A.; Bowen, R.; McMillan, I.A.; Hadfield, T.L.; Hartwig, A.; Blackburn, J.K.; Norris, M.H. Toxin and Capsule Production by *Bacillus Cereus* Biovar *Anthracis* Influence Pathogenicity in Macrophages and Animal Models. PLoS Negl Trop Dis 2024, 18, e0012779, doi:10.1371/journal.pntd.0012779.

18. Gogarten, J.F.; Düx, A.; Mubemba, B.; Pléh, K.; Hoffmann, C.; Mielke, A.; Müller-Tiburtius, J.; Sachse, A.; Wittig, R.M.; Calvignac-Spencer, S.;, et al. Tropical Rainforest Flies Carrying Pathogens Form Stable Associations with Social Nonhuman Primates. Mol Ecol 2019, 28, 4242–4258, doi:10.1111/mec.15145.

19. Jiranantasak, T.; Benn, J.S.; Metrailer, M.C.; Sawyer, S.J.; Burns, M.Q.; Bluhm, A.P.; Blackburn, J.K.; Norris, M.H. Characterization of *Bacillus Anthracis* Replication and Persistence on Environmental Substrates Associated with Wildlife Anthrax Outbreaks. PLoS One 2022, 17, e0274645, doi:10.1371/journal.pone.0274645.

20. Borst, L.; Howaldt, S.; Berdaguer, R.; Schaudinn, C.; Dupke, S.; Stämmler, M.; Lasch, P.; Scholz, H.C.; Klee, S.R. Anthrax-Causing Bacteria Form Biofilms on a Soil-Inspired Porous Glass Bead Model. PLoS Negl Trop Dis 2026, 20, e0014043, doi:10.1371/journal.pntd.0014043.

21. Turell, M.J.; Knudson, G.B. Mechanical Transmission of *Bacillus Anthracis* by Stable Flies (Stomoxys Calcitrans) and Mosquitoes (Aedes Aegypti and Aedes Taeniorhynchus). Infect Immun 1987, 55, 1859–1861, doi:10.1128/iai.55.8.1859-1861.1987.

22. Hudson, M.J.; Beyer, W.; Böhm, R.; Fasanella, A.; Garofolo, G.; Golinski, R.; Goossens, P.L.; Hahn, U.; Hallis, B.; King, A.;, et al. *Bacillus Anthracis*: Balancing Innocent Research with Dual-Use Potential. Int J Med Microbiol 2008, 298, 345–364, doi:10.1016/j.ijmm.2007.09.007.

23. Fasanella, A.; Scasciamacchia, S.; Garofolo, G.; Giangaspero, A.; Tarsitano, E.; Adone, R. Evaluation of the House Fly Musca Domestica as a Mechanical Vector for an Anthrax. PLoS One 2010, 5, e12219, doi:10.1371/journal.pone.0012219.

24. Blackburn, J.K.; Curtis, A.; Hadfield, T.L.; O’Shea, B.; Mitchell, M.A.; Hugh-Jones, M.E. Confirmation of *Bacillus Anthracis* from Flesh-Eating Flies Collected during a West Texas Anthrax Season. J Wildl Dis 2010, 46, 918–922, doi:10.7589/0090-3558-46.3.918.

25. Gainer, R.; Oksanen, A. Anthrax and the Taiga. Can Vet J 2012, 53, 1123–1125.

26. Shadomy, S.; El Idrissi, A.; Raizman, E.; Bruni, M.; Palamara, E.; Pittiglio, C.; Lubroth, J. Anthrax Outbreaks: A Warning for Improved Prevention, Control and Heightened Awareness 2016.

27. 27. Crosmary, W.; Yenamau, A.; Düx, A.; Schlotterbeck, J.; Lumbu, C.-P. Zoonosis Risks along Bushmeat Supply Chains in Central Africa. The Case of the Salonga Landscape as Source, Kinshasa and Other Major Urban Centres as Destination, Democratic Republic of Congo 2024.

28. Martínez-Campreciós, J.; Moreno, M.; Salvador, F.; Barrio-Tofiño, E.D.; Nindia, A.; Aznar, M.L.; Molina, I. Impact of Traditional Cutaneous Scarification on Anthrax Lesions: A Series of Cases from Cubal, Angola. Int J Infect Dis 2024, 140, 104–109, doi:10.1016/j.ijid.2024.01.004.

29. Kurpiers, L.A.; Schulte-Herbrüggen, B.; Ejotre, I.; Reeder, D.M. Bushmeat and Emerging Infectious Diseases: Lessons from Africa. In Problematic Wildlife; Angelici, F.M., Ed.; Springer International Publishing: Cham, 2016; pp. 507–551 ISBN 978-3-319-22245-5.

30. Pike, B.L.; LeBreton, M.; Syalors, K.; Le Doux, D.J.; Fair, J.N.; Rimoin, A.W.; Ortiz, N.; Djoko, C.F.; Tamoufe, U.; Wolfe, N. Bushmeat and Infectious Disease Emergence. In New Directions in Conservation Medicine: Applied Cases of Ecological Health; 2012; pp. 164–178.

31. Milbank, C.; Vira, B. Wildmeat Consumption and Zoonotic Spillover: Contextualising Disease Emergence and Policy Responses. Lancet Planet Health 2022, 6, e439–e448, doi:10.1016/S2542-5196(22)00064-X.

32. Kwizera, P.; Migisha, R.; Katumba, H.; Nabatta, E.; Gidudu, S.; Kwesiga, B.; Morukileng, J.; Bulage, L.; Ario, A.R. Cutaneous Anthrax Outbreak Associated with Use of Cattle Hides and Handling Carcasses, Amudat District, Uganda, 2023-2024. PLoS One 2025, 20, e0336769, doi:10.1371/journal.pone.0336769.

33. World Health Organization. Disease Outbreak News, Anthrax - Zambia, 8 december 2023. Available online: https://www.who.int/emergencies/disease-outbreak-news/item/2023-DON497 (accessed on 26 January 2026).

34. World Health Organization. Anthrax in Humans and Animals, 4th ed.; World Health Organization, Food and Agriculture Organization of the United Nations & World Organisation for Animal Health, 2008. Geneva, Switzerland. Available online: https://www.who.int/publications/i/item/9789241547536 (accessed on 1 september 2025).

35. Sprynski, N.; Valade, E.; Neulat-Ripoll, F. *Galleria Mellonella* as an Infection Model for Select Agents. In Host-Bacteria Interactions; Springer Protocols, 2014.

36. Corre, J.-P.; Piris-Gimenez, A.; Moya-Nilges, M.; Jouvion, G.; Fouet, A.; Glomski, I.J.; Mock, M.; Sirard, J.-C.; Goossens, P.L. In Vivo Germination of *Bacillus Anthracis* Spores during Murine Cutaneous Infection. J Infect Dis 2013, 207, 450–457, doi:10.1093/infdis/jis686.

37. Rougeaux, C.; Becher, F.; Ezan, E.; Tournier, J.-N.; Goossens, P.L. In Vivo Dynamics of Active Edema and Lethal Factors during Anthrax. Sci Rep 2016, 6, 23346, doi:10.1038/srep23346.

38. Rougeaux, C.; Becher, F.; Goossens, P.L.; Tournier, J.-N. Very Early Blood Diffusion of the Active Lethal and Edema Factors of *Bacillus Anthracis* After Intranasal Infection. J Infect Dis 2020, 221, 660–667, doi:10.1093/infdis/jiz497.

39. Malmquist, J.A.; Rogan, M.R.; McGillivray, S.M. *Galleria Mellonella* as an Infection Model for *Bacillus Anthracis* Sterne. Front Cell Infect Microbiol 2019, 9, 360, doi:10.3389/fcimb.2019.00360.

40. Ramarao, N.; Nielsen-Leroux, C.; Lereclus, D. The Insect *Galleria Mellonella* as a Powerful Infection Model to Investigate Bacterial Pathogenesis. J Vis Exp 2012, e4392, doi:10.3791/4392.

41. Lowe, D.E.; Ya, J.; Glomski, I.J. In Trans Complementation of Lethal Factor Reveal Roles in Colonization and Dissemination in a Murine Mouse Model. PLoS One 2014, 9, e95950, doi:10.1371/journal.pone.0095950.

42. Weiner, Z.P.; Boyer, A.E.; Gallegos-Candela, M.; Cardani, A.N.; Barr, J.R.; Glomski, I.J. Debridement Increases Survival in a Mouse Model of Subcutaneous Anthrax. PLoS One 2012, 7, e30201, doi:10.1371/journal.pone.0030201.

43. Glomski, I.J.; Piris-Gimenez, A.; Huerre, M.; Mock, M.; Goossens, P.L. Primary Involvement of Pharynx and Peyer’s Patch in Inhalational and Intestinal Anthrax. PLoS Pathog 2007, 3, e76, doi:10.1371/journal.ppat.0030076.

44. Bischof, T.S.; Hahn, B.L.; Sohnle, P.G. Characteristics of Spore Germination in a Mouse Model of Cutaneous Anthrax. J Infect Dis 2007, 195, 888–894, doi:10.1086/511824.

45. Sirard, J.C.; Guidi-Rontani, C.; Fouet, A.; Mock, M. Characterization of a Plasmid Region Involved in *Bacillus Anthracis* Toxin Production and Pathogenesis. Int J Med Microbiol 2000, 290, 313–316, doi:10.1016/S1438-4221(00)80030-2.

46. Cote, C.K.; Welkos, S.L. Anthrax Toxins in Context of *Bacillus Anthracis* Spores and Spore Germination. Toxins (Basel) 2015, 7, 3167–3178, doi:10.3390/toxins7083167.

47. Boyer, A.E.; Gallegos-Candela, M.; Lins, R.C.; Solano, M.I.; Woolfitt, A.R.; Lee, J.S.; Sanford, D.C.; Knostman, K.A.B.; Quinn, C.P.; Hoffmaster, A.R.;, et al. Comprehensive Characterization of Toxins during Progression of Inhalation Anthrax in a Non-Human Primate Model. PLoS Pathog 2022, 18, e1010735, doi:10.1371/journal.ppat.1010735.

48. Lins, R.C.; Boyer, A.E.; Kuklenyik, Z.; Woolfitt, A.R.; Goldstein, J.; Hoffmaster, A.R.; Gallegos-Candela, M.; Leysath, C.E.; Chen, Z.; Brumlow, J.O.;, et al. Zeptomole per Milliliter Detection and Quantification of Edema Factor in Plasma by LC-MS/MS Yields Insights into Toxemia and the Progression of Inhalation Anthrax. Anal Bioanal Chem 2019, 411, 2493–2509, doi:10.1007/s00216-019-01730-4.

49. Dumetz, F.; Jouvion, G.; Khun, H.; Glomski, I.J.; Corre, J.-P.; Rougeaux, C.; Tang, W.-J.; Mock, M.; Huerre, M.; Goossens, P.L. Noninvasive Imaging Technologies Reveal Edema Toxin as a Key Virulence Factor in Anthrax. Am J Pathol 2011, 178, 2523–2535, doi:10.1016/j.ajpath.2011.02.027.

50. Hurst, J.R.; Shannon, B.A.; Craig, H.C.; Rishi, A.; Tuffs, S.W.; McCormick, J.K. The *Streptococcus Pyogenes* Hyaluronic Acid Capsule Promotes Experimental Nasal and Skin Infection by Preventing Neutrophil-Mediated Clearance. PLoS Pathog 2022, 18, e1011013, doi:10.1371/journal.ppat.1011013.

51. Wessels, M.R.; Goldberg, J.B.; Moses, A.E.; DiCesare, T.J. Effects on Virulence of Mutations in a Locus Essential for Hyaluronic Acid Capsule Expression in Group A Streptococci. Infect Immun 1994, 62, 433–441, doi:10.1128/iai.62.2.433-441.1994.

52. Dale, J.B.; Washburn, R.G.; Marques, M.B.; Wessels, M.R. Hyaluronate Capsule and Surface M Protein in Resistance to Opsonization of Group A Streptococci. Infect Immun 1996, 64, 1495–1501, doi:10.1128/iai.64.5.1495-1501.1996.

53. Schrager, H.M.; Rheinwald, J.G.; Wessels, M.R. Hyaluronic Acid Capsule and the Role of Streptococcal Entry into Keratinocytes in Invasive Skin Infection. J Clin Invest 1996, 98, 1954–1958, doi:10.1172/JCI118998.

54. Russell, B.H.; Liu, Q.; Jenkins, S.A.; Tuvim, M.J.; Dickey, B.F.; Xu, Y. In Vivo Demonstration and Quantification of Intracellular *Bacillus Anthracis* in Lung Epithelial Cells. Infect Immun 2008, 76, 3975–3983, doi:10.1128/IAI.00282-08.

55. van Sorge, N.M.; Ebrahimi, C.M.; McGillivray, S.M.; Quach, D.; Sabet, M.; Guiney, D.G.; Doran, K.S. Anthrax Toxins Inhibit Neutrophil Signaling Pathways in Brain Endothelium and Contribute to the Pathogenesis of Meningitis. PLoS One 2008, 3, e2964, doi:10.1371/journal.pone.0002964.

56. Russell, B.H.; Vasan, R.; Keene, D.R.; Xu, Y. *Bacillus Anthracis* Internalization by Human Fibroblasts and Epithelial Cells. Cell Microbiol 2007, 9, 1262–1274, doi:10.1111/j.1462-5822.2006.00869.x.

57. Tournier, J.-N.; Rossi Paccani, S.; Quesnel-Hellmann, A.; Baldari, C.T. Anthrax Toxins: A Weapon to Systematically Dismantle the Host Immune Defenses. Mol Aspects Med 2009, 30, 456–466, doi:10.1016/j.mam.2009.06.002.

58. Paccani, S.R.; Baldari, C.T. T Cell Targeting by Anthrax Toxins: Two Faces of the Same Coin. Toxins (Basel) 2011, 3, 660–671, doi:10.3390/toxins3060660.

59. Weiner, Z.P.; Glomski, I.J. Updating Perspectives on the Initiation of *Bacillus Anthracis* Growth and Dissemination through Its Host. Infect Immun 2012, 80, 1626–1633, doi:10.1128/IAI.06061-11.

60. Lynskey, N.N.; Banerji, S.; Johnson, L.A.; Holder, K.A.; Reglinski, M.; Wing, P.A.C.; Rigby, D.; Jackson, D.G.; Sriskandan, S. Rapid Lymphatic Dissemination of Encapsulated Group A Streptococci via Lymphatic Vessel Endothelial Receptor-1 Interaction. PLoS Pathog 2015, 11, e1005137, doi:10.1371/journal.ppat.1005137.

61. Moayeri, M.; Haines, D.; Young, H.A.; Leppla, S.H. *Bacillus Anthracis* Lethal Toxin Induces TNF-α–Independent Hypoxia-Mediated Toxicity in Mice. J. Clin. Invest. 2003, 112, 670–682, doi:10.1172/JCI17991.

62. Liu, S.; Moayeri, M.; Leppla, S.H. Anthrax Lethal and Edema Toxins in Anthrax Pathogenesis. Trends Microbiol 2014, 22, 317–325, doi:10.1016/j.tim.2014.02.012.

63. Izbanova, U.; Duisenova, A.; Tokmurziyeva, G.; Abdrakhmanova, A.; Aibosynova, S.; Kosherova, B.; Yegemberdiyeva, R.; Sadykova, A.; Umarova, S.; Askarov, D.;, et al. Severe Cutaneous Anthrax with Systemic Complications: A Case Report. Front. Med. 2026, 13, 1804212, doi:10.3389/fmed.2026.1804212.

64. Yuan, H.; Zheng, Y.; Zhang, W.; Xie, H. Two Cases of Human Cutaneous Anthrax with Massive Tissue Damage, Severe Edema, and Slight Injury to the Liver. Int J Dermatology 2018, 57, 358–361, doi:10.1111/ijd.13865.

65. Firoved, A.M.; Miller, G.F.; Moayeri, M.; Kakkar, R.; Shen, Y.; Wiggins, J.F.; McNally, E.M.; Tang, W.-J.; Leppla, S.H. *Bacillus Anthracis* Edema Toxin Causes Extensive Tissue Lesions and Rapid Lethality in Mice. The American Journal of Pathology 2005, 167, 1309–1320, doi:10.1016/S0002-9440(10)61218-7.

66. Watson, L.E.; Kuo, S.; Katki, K.; Dang, T.; Park, S.K.; Dostal, D.E.; Tang, W.-J.; Leppla, S.H.; Frankel, A.E. Anthrax Toxins Induce Shock in Rats by Depressed Cardiac Ventricular Function. PLoS ONE 2007, 2, e466, doi:10.1371/journal.pone.0000466.

67. Moayeri, M.; Crown, D.; Dorward, D.W.; Gardner, D.; Ward, J.M.; Li, Y.; Cui, X.; Eichacker, P.; Leppla, S.H. The Heart Is an Early Target of Anthrax Lethal Toxin in Mice: A Protective Role for Neuronal Nitric Oxide Synthase (nNOS). PLoS Pathog 2009, 5, e1000456, doi:10.1371/journal.ppat.1000456.

68. Liu, S.; Zhang, Y.; Moayeri, M.; Liu, J.; Crown, D.; Fattah, R.J.; Wein, A.N.; Yu, Z.-X.; Finkel, T.; Leppla, S.H. Key Tissue Targets Responsible for Anthrax-Toxin-Induced Lethality. Nature 2013, 501, 63–68, doi:10.1038/nature12510.

69. Kuo, S.-R.; Willingham, M.C.; Bour, S.H.; Andreas, E.A.; Park, S.K.; Jackson, C.; Duesbery, N.S.; Leppla, S.H.; Tang, W.-J.; Frankel, A.E. Anthrax Toxin-Induced Shock in Rats Is Associated with Pulmonary Edema and Hemorrhage. Microbial Pathogenesis 2008, 44, 467–472, doi:10.1016/j.micpath.2007.12.001.

70. Akdeniz, N.; Calka, O.; Ozkol, H.U.; Akdeniz, H. Cutaneous Anthrax Resulting in Renal Failure with Generalized Tissue Damage. Cutaneous and Ocular Toxicology 2013, 32, 327–329, doi:10.3109/15569527.2013.768257.

71. Hicks, C.W.; Cui, X.; Sweeney, D.A.; Li, Y.; Barochia, A.; Eichacker, P.Q. The Potential Contributions of Lethal and Edema Toxins to the Pathogenesis of Anthrax Associated Shock. Toxins 2011, 3, 1185–1202, doi:10.3390/toxins3091185.

72. Hendricks, K.; Martines, R.B.; Bielamowicz, H.; Boyer, A.E.; Long, S.; Byers, P.; Stoddard, R.A.; Taylor, K.; Kolton, C.B.; Gallegos-Candela, M.;, et al. Welder’s Anthrax: A Tale of 2 Cases. Clin Infect Dis 2022, 75, S354–S363, doi:10.1093/cid/ciac535.

73. Leendertz, F.H.; Lankester, F.; Guislain, P.; Néel, C.; Drori, O.; Dupain, J.; Speede, S.; Reed, P.; Wolfe, N.; Loul, S.;, et al. Anthrax in Western and Central African Great Apes. Am J Primatol 2006, 68, 928–933, doi:10.1002/ajp.20298.

